# E3 Ubiquitin Ligase CHIP Drives Monoubiquitination-mediated Nuclear Import of Tumor Suppressor PTEN

**DOI:** 10.1101/2023.01.02.522453

**Authors:** Shrabastee Chakraborty, Subhajit Karmakar, Malini Basu, Mrinal K. Ghosh

## Abstract

Nuclear localization of Phosphatase and Tensin Homolog Deleted on Chromosome TEN (PTEN) is crucial for its tumor suppressive activities. Monoubiquitination has been established as a principal mechanism driving nuclear translocation of PTEN. In this study, we describe a novel mechanism wherein Carboxy terminus of Hsc70 Interacting Protein (CHIP) mediates monoubiquitination of PTEN, leading to the latter’s nuclear import. Analysis of subcellular fractions of multiple cell lines revealed a rise in both nuclear and total cellular PTEN levels under monoubiquitination-promoting conditions, an effect that was abrogated when CHIP expression was silenced. We established a time-point kinetics of CHIP-mediated nuclear translocation of PTEN using immunocytochemistry. We identified a role of Karyopherin α1 (KPNA1) in facilitating nuclear transport of monoubiquitinated PTEN. We further established a direct interaction between CHIP and PTEN inside the nuclear compartment, where CHIP could participate in either polyubiquitination or monoubiquitination of the latter. CHIP was found to monoubiquitinate PTEN on K13 and K289 residues. Finally, we showed that CHIP-mediated monoubiquitination of PTEN resulted in increased apoptosis, decreased cell viability and proliferation. This enhancement of PTEN’s tumor suppressive abilities by CHIP involved modulation of transcriptional activities of PTEN. To the best of our knowledge, this is the first report elucidating CHIP’s non-canonical roles on PTEN, which is also established here as a nuclear interacting partner of CHIP.

## Introduction

PTEN is a prominent tumor suppressor that has been dubbed “a new guardian of genome”.^1^ Although predominantly cytoplasmic, PTEN localization in nucleus, endoplasmic reticulum, and mitochondria has been observed. Among these, nuclear localization of PTEN has gained importance as a vast repertoire of anti-tumorigenic functions exclusive to nuclear PTEN has been unearthed. Nuclear PTEN participates in DNA damage response, genomic stability maintenance, replication fork maintenance, cell cycle regulation, transcriptional and post-transcriptional modulation, and apoptosis.^2^

Lacking a classic Nuclear Localization Signal (NLS),^3,4^ PTEN’s nuclear translocation relies on passive diffusion,^5^ Ran-mediated transport,^6^ major vault protein-mediated transport,^3,7^ phosphorylation,^8,9^ SUMOylation,^10^ NEDDylation,^11^ and finally, monoubiquitination. NEDD4-1, the first E3 ligase known to cause polyubiquitination-mediated proteasomal degradation of PTEN,^12^ was also established as a monoubiquitinating E3 ligase that promoted its nuclear localization.^13^ Subsequently, XIAP was identified as an E3 ligase capable of both poly- and monoubiquitinating PTEN, leading to degradation and nuclear import, respectively.^14^ However, these E3 ligases primarily target cytoplasmic PTEN, and reports on ubiquitin-mediated regulation of nuclear PTEN are lacking. To date, a single E3 ligase, FBXO22, has been shown to selectively degrade nuclear PTEN *via* ubiquitin-proteasome pathway, with negligible effect on cytoplasmic or total cellular PTEN.^15^ Further, the exact mechanism through which monoubiquitinated PTEN enters the nucleus remains largely unexplored, although possible roles of Importin11 have been described.^16^

E3 ligase CHIP can promote non-canonical ubiquitination of its substrates, including monoubiquitination.^17^ It can also chaperone its substrates inside the nucleus^18^ and perform E3 ligase activities within the nuclear compartment.^19^ We have previously shown that CHIP mediates polyubiquitination and proteasomal degradation of PTEN.^20^ However, CHIP overexpression revealed both poly- and monoubiquitinated PTEN adducts while CHIP knockdown resulted in reduced level of nuclear PTEN. Hence, we envisioned a probable role of CHIP in driving monoubiquitination-mediated nuclear import of PTEN.

In this study, we reported a non-canonical function of CHIP where it monoubiquitinates PTEN, causing its nuclear entry. We further identified CHIP as an interacting partner and E3 ligase of PTEN inside the nuclear compartment. In addition, we demonstrated the involvement of KPNA1 protein in enhancing nuclear translocation of monoubiquitinated PTEN.

## Materials and Methods

### Bioinformatic analyses

PTEN gene expression was analyzed in 22 cancers from The Cancer Genome Atlas (TCGA) dataset using TNM plot.^21^ PTEN mRNA expression was analyzed in colon adenocarcinoma, liver hepatocellular carcinoma, and cervical squamous cell and endocervical adenocarcinoma using Gene Expression Profiling Interactive Analysis (GEPIA).^22^ UALCAN (http://ualcan.path.uab.edu/analysis.html) analysis of PTEN protein expression was performed in colon and liver cancer using Clinical Proteomic Tumor Analysis Consortium (CPTAC) data.^23^ The specimens were compared as cancer *vs* normal patients. *P <* 0.01 was considered significant. Median mRNA and protein expressions of PTEN were explored in HEK293, HCT 116, HepG2, and HeLa cell lines using ProteomicsDB.^24^ cBioPortal^25^ was used to analyze the correlations between PTEN and its downstream targets *viz*., Rad51, Dre1, and VEGF in colon, liver, and cervical cancers using TCGA PanCancer dataset and Cancer Cell Line Encyclopedia. *P <* 0.05 was considered significant.

### Cell culture and transfection

HCT 116 cells were cultured in McCoy’s 5A medium, HEK293, HepG2, and HeLa cells in Dulbecco’s Modified Eagle’s Medium (DMEM) with 10% heat-inactivated Fetal Bovine Serum and antibiotics Penicillin, Streptomycin, and Gentamycin at recommended doses. Cells were maintained at 37°C with 5% CO_2_. DNA constructs were transfected using PEI (polyethylenimine, Sigma Aldrich) in HEK293, and Lipofectamine 3000 (Invitrogen) in other cancer cell lines. siRNAs were transfected using Lipofectamine RNAimax (Invitrogen).

### Cloning and Plasmid constructs

pGZ21dx-GFP-PTEN-WT (GFP-PTEN), pIRES-CHIP, pIRES-CHIP-H260Q, pIRES-CHIP-K30A, pGZ21dx-GFP-XIAP, FLAG-USP7 plasmids were described previously.^20,26^ PRK5-HA-Ubiquitin-WT, pRK5-HA-Ubiquitin-KØ, pRK5-HA-Ubiquitin-K48R, pCI-HA-Nedd4-1, and HA-NLS-PTEN were procured from Addgene. NLS-CHIP was generated by replacing EBFP from pEBFP-nuc (Addgene, 14893) with CHIP insert excised from pIRES-CHIP.

### Short interfering RNA (siRNA) and short hairpin RNA (shRNA) mediated knockdown

Control shRNA and shRNA against PTEN were purchased from Addgene (86645), Scramble siRNA, CHIP siRNA (sc-43555), and USP7 siRNA (sc-41521), purchased from Santa Cruz Biotechnology, were added at a final concentration of 30 nM.

### Site directed mutagenesis

PTEN monoubiquitination mutants PTEN-K13R and PTEN-K289R were generated from GFP-PTEN by site directed mutagenesis using QuickChange XL Site Directed Mutagenesis Kit (Agilent technologies). Table S1 includes the primer sequences.

### Antibodies and chemicals

The primary antibodies used were PTEN, CHIP, GAPDH, Lamin B, GFP, ubiquitin, USP7, cleaved PARP (Santa Cruz Biotechnolgy); PTEN, cleaved Caspase 3, Bax, Bim (Cell Signaling Technology); CHIP (AbCam); Actin, HA-tag, FLAG-tag, (Sigma Aldrich); NEDD4-1 (Abclonal). HRP-tagged anti-goat, anti-mouse, anti-rabbit secondary antibodies, Ivermectin (I8898, diluted in 95% ethanol), and Leptomycin B (L2913, diluted in 100% ethanol) were purchased from Sigma Aldrich.

### Whole cell lysate preparation, subcellular fractionation, and Western blotting

Whole cell lysate preparation, nucleo-cytoplasmic fractionation, and western blotting were performed as described previously.^27^ Blots were developed with Millipore Luminata Classico in ChemiDoc MP Imaging System (Biorad). Densitometric quantification was done using ImageJ. Loading controls for whole cell lysate (WCL), nuclear extract (NE), and cytoplasmic extract (CE) were Actin, Lamin B, and GAPDH, respectively.

### Co-immunoprecipitation

Co-immunoprecipitation was performed in HEK293 cells as described previously.^26^ Nuclear lysate (500 µg) or total protein (1 mg) was used for immunoprecipitation.

### Immunofluorescence microscopy

Immunofluorescence microscopy was performed in HEK293 cells as described previously.^27^ Primary antibodies against PTEN, KPNA1 (Santa Cruz Biotechnologies), CHIP (AbCam), and fluorescence-conjugated secondary antibodies (Alexafluor488, Alexafluor594) were used. The nuclei were stained with Hoechst 33342 (Life Technologies). The images were captured at 60x in Zeiss LSM 980 confocal microscope using ZENblue3 software and analyzed using ImageJ.

### cDNA preparation and quantitative PCR

TRIZOL (Invitrogen) was used to isolate total RNA as per the manufacturer’s protocol. 2 μg of RNA was converted to cDNA using PrimeScript 1^st^ Strand cDNA Synthesis Kit (TaKaRa) in SureCycler 8800 (Agilent Technologies). Quantitative Real time PCR was performed using FastStart universal SyBR Green Master Mix (Roche) in 7500 Fast Real-time PCR System (Applied Biosystems). The Standard deviation calculation and quantification were performed from three independent experiments using 18s rRNA as the internal control. Table S1 includes the primer sequences.

### Cell viability assay

Cell viability assay was performed using MTT [3-(4,5-dimethylthiazol-2-yl)-2,5-diphenyl tetrazolium bromide] on HEK293 and HCT 116 cells 48 hrs post-transfection, as described previously.

### Wound healing (scratch) assay

Wound healing assay was performed in HEK293, HCT 116, HepG2, and HeLa cells as described previously.^27^ Images were captured at 0 hr and 48 hrs post-transfection.

### Colony formation assay

Colony formation assay was performed in HEK293, HCT 116, HepG2, and HeLa cells after transfection, as described previously.^27^

### Statistical analysis

The statistical significance was drawn by conducting Two Samples Student’s *t*-test, with significance values represented as * (*P* ≤ 0.05), ** (*P* ≤ 0.01), *** (*P* ≤ 0.001). The statistical analyses were performed using GraphPad Prism 8 (https://www.graphpad.com/scientific-software/prism).

## Results

### CHIP-mediated non-canonical ubiquitination increases cellular PTEN pool

Although nuclear PTEN is primarily tumor-suppressive, its levels are variable in different cancers. For our study, we selected colorectal, hepatic, and cervical cancers as they exhibit differential subcellular distribution patterns of PTEN. In colorectal cancer, nuclear PTEN loss was positively correlated with disease progression.^28–32^ In invasive cervical carcinoma, nuclear egress of PTEN was observed^33^ although predominantly nuclear PTEN expression was reported in normal and malignant endocervical cells.^34^ While both cytoplasmic and nuclear PTEN decreased in tumor tissues of hepatic cancers,^35^ another study found nuclear PTEN in tumor cells and cytoplasmic PTEN in surrounding cells.^36^

Using TNM plot, a comparative analysis of PTEN gene expression of normal and tumor tissues in 22 different cancers was performed with RNAseq data from TCGA database,^21^ which revealed significant differences in colon cancer but not in liver cancer (Fig. 1A). PTEN mRNA levels in colon adenocarcinoma, liver hepatocellular carcinoma, and cervical squamous cell and endocervical adenocarcinoma were further assessed in normal *vs* tumor samples using GEPIA^22^ (Fig. 1B). UALCAN analysis in normal *vs* tumor tissues of colon and liver cancer samples from CPTAC datasets^23^ showed that PTEN protein expression was significantly lower in tumor samples in both cancers (Fig. 1C). For our study, we chose HCT 116, HepG2, and HeLa cell lines, belonging to colon, hepatic, and cervical cancers, respectively, while exploring mechanistic details in HEK293. PTEN mRNA and protein expressions in HCT 116, HepG2, HeLa, and HEK293 were assessed using quantitative mass spectrometry-based proteomics data from ProteomicsDB^24^ (Fig. 1D). Total cellular and subcellular (cytoplasmic and nuclear) distribution of endogenous PTEN and CHIP were checked in HCT 116, HepG2, and HeLa (Fig. 1E).

**Figure 1.**
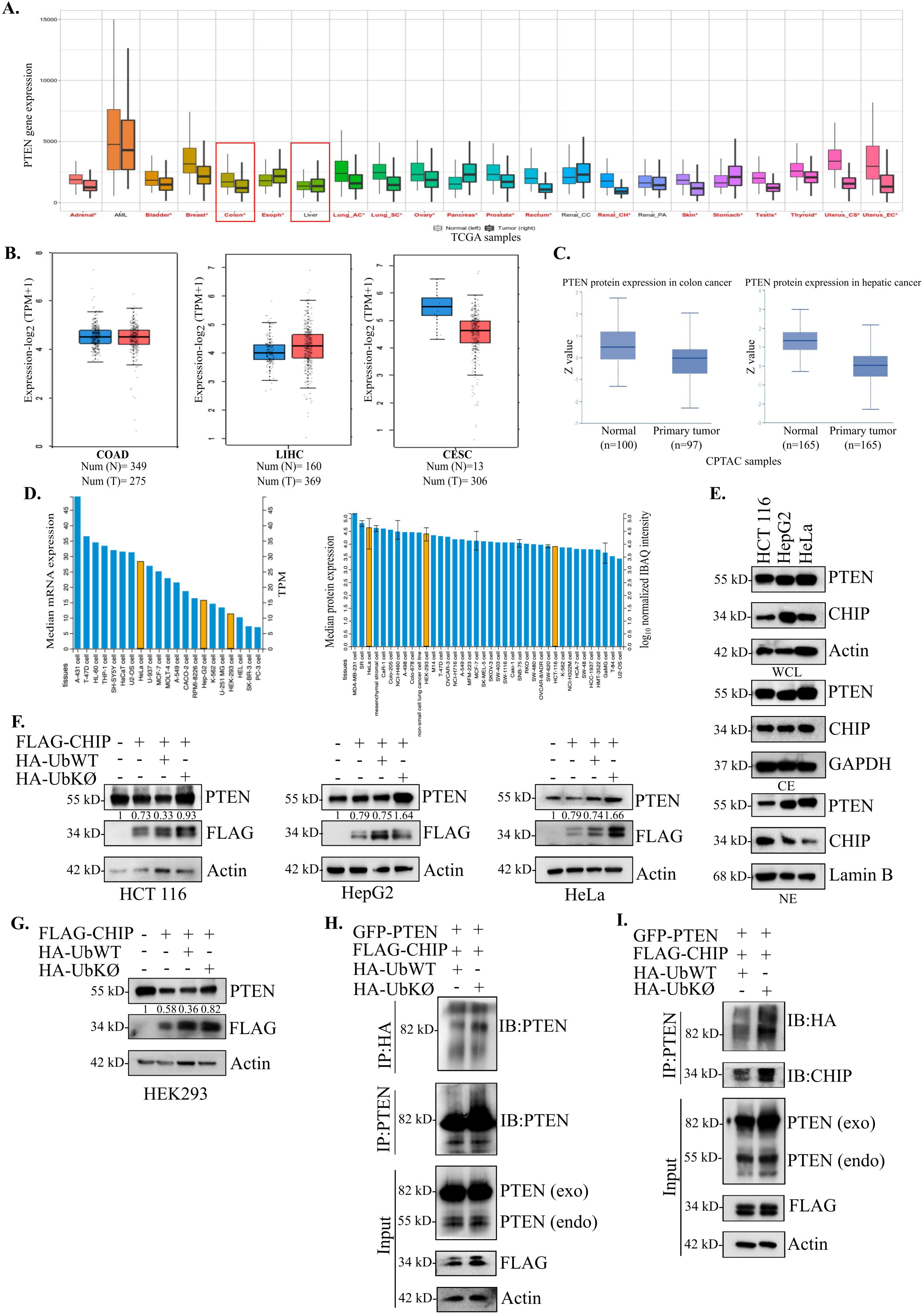
CHIP-mediated non-canonical ubiquitination increases cellular PTEN pool. (**A**) Boxplots represent differential gene expression analysis of PTEN in normal *vs* tumor tissues in 22 cancers using TNM plot from TCGA database. The cancers marked in red show significant differences in Mann-Whitney U test (*P* < 0.01). (**B**) Boxplots represent PTEN transcriptional levels (expressed as log_2_(TPM+1)) in normal *vs* tumor samples of colon adenocarcinoma (COAD), liver hepatocellular carcinoma (LIHC), and cervical squamous cell and endocervical adenocarcinoma (CESC) using GEPIA. (**C**) Boxplots represent UALCAN analysis of PTEN protein levels (expressed as Z values) in normal *vs* tumor samples of colon and hepatic cancer, data obtained from CPTAC (*P* < 0.01). (**D**) Bar diagrams represent median mRNA and protein expressions of PTEN in multiple cell lines, data obtained from ProteomicsDB. Highlighted bars show HeLa, HepG2, HEK293 (left panel) and HeLa, HEK293, and HCT 116 (right panel). (**E**) Whole cell lysates, cytoplasmic, and nuclear extracts prepared from HCT 116, HepG2, and HeLa cells were immunoblotted with anti-PTEN and anti-CHIP antibodies. (**F**) FLAG-CHIP was transfected either alone or together with HA-UbWT or HA-UbKØ in HCT 116 (left panel), HepG2 (middle panel), and HeLa (right panel) cells. Whole cell lysates were probed for PTEN and FLAG (exogenous CHIP). (**G**) FLAG-CHIP was transfected either alone or together with HA-UbWT or HA-UbKØ in HEK293 cells. Whole cell lysates were probed for PTEN and FLAG (exogenous CHIP). (**H**) HEK293 cells were co-transfected with GFP-PTEN, FLAG-CHIP, and either HA-UbWT or HA-UbKØ. Whole cell lysates were immunoprecipitated using anti-HA or anti-PTEN antibodies and immunoblotted for PTEN. (**I**) HEK293 cells were co-transfected with GFP-PTEN, FLAG-CHIP, and either HA-UbWT or HA-UbKØ. Whole cell lysates were immunoprecipitated using anti-PTEN antibody and immunoblotted for HA (exogenous ubiquitin) and CHIP. 1 µg of each DNA construct was used in transfection for overexpression of cloned gene.

To detect possible modes of ubiquitination, we used wild type ubiquitin (UbWT) and its lysine-deficient mutant (UbKØ), respectively. Unlike UbWT, UbKØ was incapable of ubiquitin chain elongation with all seven lysine residues mutated to arginine, rendering the substrate molecule in monoubiquitinated and multi-monoubiquitinated states. In HCT 116, HepG2, and HeLa cell lines, the total cellular pool of PTEN decreased upon CHIP overexpression and was further reduced when UbWT was co-expressed, confirming the E3 ligase’s canonical role in PTEN degradation. However, co-expression of UbKØ restored cellular PTEN level, indicating a non-canonical monoubiquitination that did not cause its proteasomal degradation (Fig. 1F). HEK293 cells gave similar results (Fig. 1G).

After overexpressing PTEN and CHIP with either HA-UbWT or HA-UbKØ, we detected PTEN among the immunoprecipitated proteins conjugated with exogenous UbWT or UbKØ. The similar pattern of Ub-adduct formation upon using either UbWT or UbKØ suggested that CHIP, in addition to polyubiquitination, participated in multi-monoubiquitination of PTEN (Fig. 1H). In the reverse experiment, immunoprecipitation using anti-PTEN antibody revealed the existence of HA-tagged Ub-adducts (Fig. 1I). In both cases, the greater amount of PTEN pulled down in the UbKØ lane further supported the monoubiquitination mechanism that did not lead to proteasomal degradation.

### CHIP-mediated monoubiquitination leads to nuclear import of PTEN

Monoubiquitination is a bona fide mode for PTEN’s nuclear import. To see whether CHIP-mediated monoubiquitination also leads to the same fate, we performed subcellular fractionation assays from HEK293 cells with or without CHIP overexpression. Nuclear PTEN showed very little decrease while the reduction in cytoplasmic PTEN pool was more prominent (Fig. 2A), suggesting that CHIP’s effects on nuclear and cytoplasmic PTEN might be different. When CHIP was co-expressed with UbKØ in HEK293 (Fig. 2B) and HCT 116 (Fig. S1A) cells, nuclear PTEN was elevated. Exogenous PTEN level also increased in the nucleus (Fig. S1B). We conjectured CHIP-mediated monoubiquitination of PTEN to be the cause for such observations.

**Figure 2.**
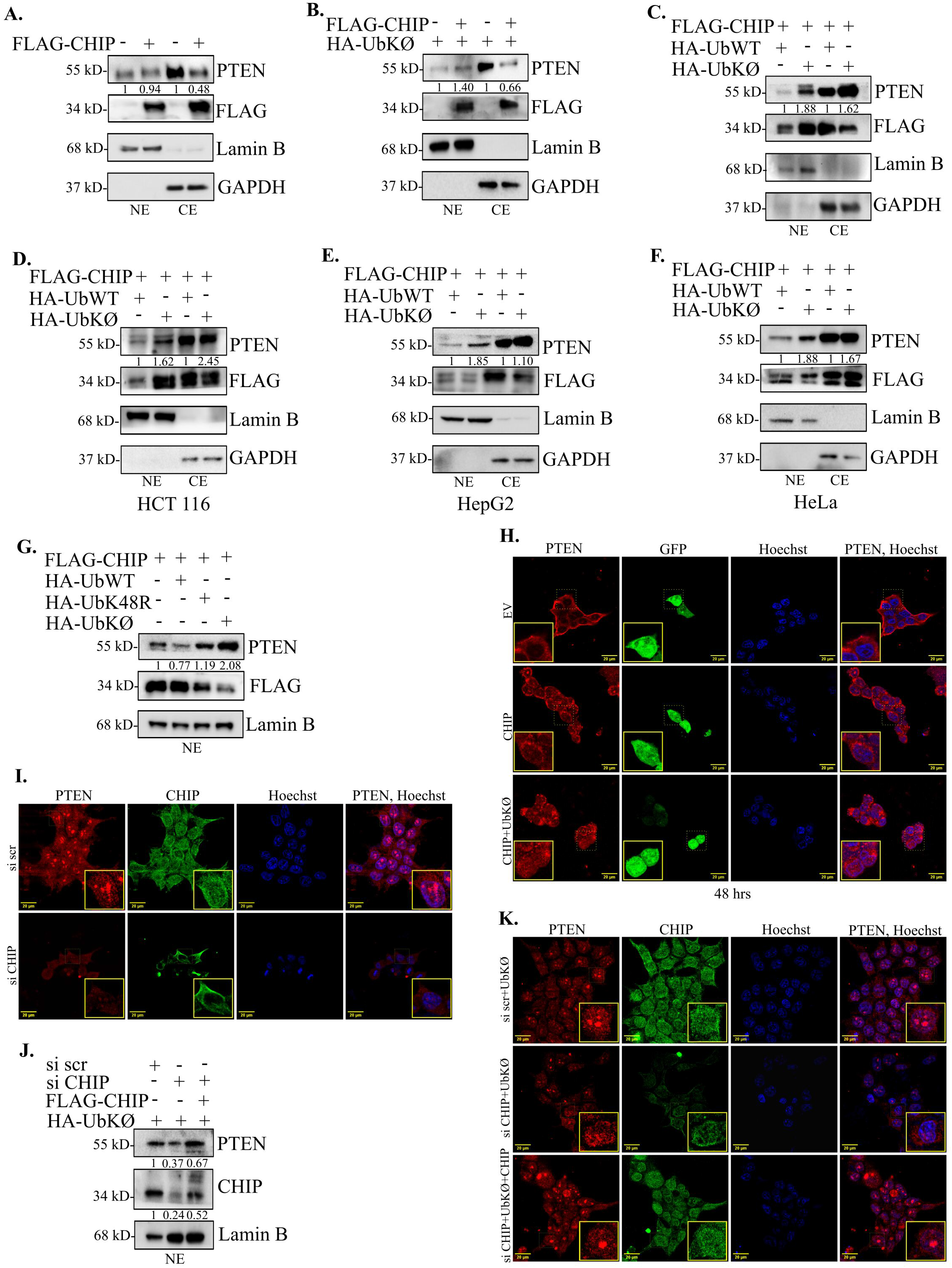
CHIP-mediated monoubiquitination leads to nuclear import of PTEN. (**A**) HEK293 cells were transfected with empty vector or FLAG-CHIP. Upon subcellular fractionation, nuclear and cytoplasmic extracts were probed for PTEN and FLAG (exogenous CHIP). (**B**) HEK293 cells were transfected with HA-UbKØ with or without FLAG-CHIP. Nuclear and cytoplasmic extracts were probed for PTEN and FLAG (exogenous CHIP). (**C-F**) FLAG-CHIP was transfected with either HA-UbWT or HA-UbKØ in HEK293 (**C**), HCT 116 (**D**), HepG2 (**E**), and HeLa (**F**) cells. Cytoplasmic and nuclear extracts were immunoblotted for PTEN and FLAG (exogenous CHIP). (**G**) HEK293 cells were transfected with FLAG-CHIP, either alone or with HA-UbWT, HA-UbK48R, or HA-UbKØ. Nuclear extracts were probed for PTEN and FLAG (exogenous CHIP). (**H**) HEK293 cells were transfected with either empty vector (EV), or FLAG-CHIP, or FLAG-CHIP and HA-UbKØ, EV and FLAG-CHIP GFP-tagged. 48 hrs post-transfection, cells were fixed and stained with primary antibodies against PTEN, secondary antibodies conjugated to AF594 (PTEN, red). GFP was observed to determine EV or CHIP overexpression. (**I**) HEK293 cells were transfected with either scrambled siRNA or CHIP siRNA (30 nM). Cells were stained with primary antibodies against PTEN and CHIP, secondary antibodies conjugated to AF594 (PTEN, red) or AF488 (CHIP, green). (**J**) HEK293 cells were transfected with either scrambled siRNA or CHIP siRNA (30 nM) in combination with either HA-UbKØ or HA-UbKØ and FLAG-CHIP. Nuclear extracts were probed for PTEN and CHIP. (**K**) HEK293 cells were transfected with scrambled siRNA or CHIP siRNA (30 nM) with either HA-UbKØ or HA-UbKØ and pCS2-HA-CHIP. 1 µg of each DNA construct was used in transfection for overexpression of cloned gene. For immunofluorescence microscopy, 60X magnification was used. Nuclei were visualized through Hoechst staining.

To elucidate the effects of poly-*vs* monoubiquitination, we co-expressed CHIP with either UbWT or UbKØ in HEK293, HCT 116, HepG2, and HeLa cell lines. Both nuclear and cytoplasmic PTEN levels showed significant increase in the case of monoubiquitination-enhancing UbKØ overexpression (Fig. 2C-F). Exogenous PTEN showed similar effects (Fig. S1C). This concomitant increase could explain the increase in overall cellular PTEN in UbKØ-expressing condition observed in Fig. 1F-G. Next, we co-expressed CHIP either alone or with UbWT, UbK48R, and UbKØ as depicted in the figure. UbK48R, whose lack of K48 residue blocked K48-mediated polyubiquitin chain elongation and subsequent degradation, drove nuclear entry of non-degraded PTEN *via* possible passive diffusion. However, UbKØ was able to further enhance the nuclear accumulation, highlighting the importance of active monoubiquitination (Fig. 2G).

To explore the kinetics of nuclear translocation of PTEN following its monoubiquitination by CHIP, HEK293 cells transfected with either empty vector, CHIP, or CHIP supplemented with UbKØ, were fixed at the designated time-points before performing immunocytochemistry. Up to 24 hrs, PTEN subcellular distribution did not differ much under these three experimental conditions. From the 30 hrs time-point, however, CHIP expression led to more nuclear localization of PTEN, which increased further when UbKØ was co-expressed (Fig. S1D, upper panel). In the subsequent time-points, more PTEN gradually localized to the nucleus, an effect that was the most pronounced under UbKØ co-expression (Fig. S1D, middle and lower panel). At 48 hrs post-transfection, control cells showed predominantly cytoplasmic PTEN, CHIP-transfected cells exhibited both nuclear and cytoplasmic PTEN, although cytoplasmic levels were considerably higher than nuclear. In contrast, upon co-transfecting CHIP and UbKØ, PTEN exhibited more nuclear distribution (Fig. 2H).

Although siRNA-mediated CHIP depletion could increase cytoplasmic PTEN level, surprisingly, it diminished the nuclear PTEN pool.^20^ Confocal imaging showed a prominent nuclear egress of PTEN when CHIP was knocked down in HEK293 (Fig. 2I) and HeLa cells,^20^ further corroborating its role in maintaining nuclear PTEN. Even enhancing monoubiquitination by overexpressing UbKØ could not increase nuclear PTEN under CHIP-depleted condition, while reintroduction of CHIP restored it (Fig. 2J). Immunofluorescence experiments also revealed similar rescue effects when CHIP was reintroduced (Fig. 2K). Taken together, CHIP could drive both poly- and monoubiquitination of PTEN, with the latter causing PTEN’s nuclear import.

### Nucleo-cytoplasmic shuttling of monoubiquitinated PTEN is dependent on nuclear import/export machinery

To explore the mechanism by which monoubiquitinated PTEN gains nuclear entry, we transfected HEK293 cells with UbKØ with or without CHIP overexpression, and treated with increasing doses of Ivermectin, a karyopherin inhibitor. Western blot analysis of the nuclear fraction revealed that Ivermectin inhibited nuclear import of monoubiquitinated PTEN in a dose-dependent manner (Fig. 3A). To determine the possible involvement of various karyopherins, we co-expressed Karyopherin α1 (KPNA1), Karyopherin α2 (KPNA2), and Karyopherin α4 (KPNA4) with CHIP and UbKØ. KPNA1 co-expression further enhanced nuclear PTEN, suggesting its involvement in monoubiquitinated PTEN’s nuclear import (Fig. 3B). An immunoprecipitation experiment using the nuclear fraction indicated monoubiquitinated PTEN pull-down under KPNA1 overexpression (Fig. 3C). Immunofluorescence studies on HEK293 cells transfected with either empty vector, CHIP, or CHIP and UbKØ revealed that PTEN-KPNA1 colocalization was the most prominent under UbKØ co-expression. As expected, this condition resulted in the highest level of nuclear PTEN (Fig. 3D).

**Figure 3.**
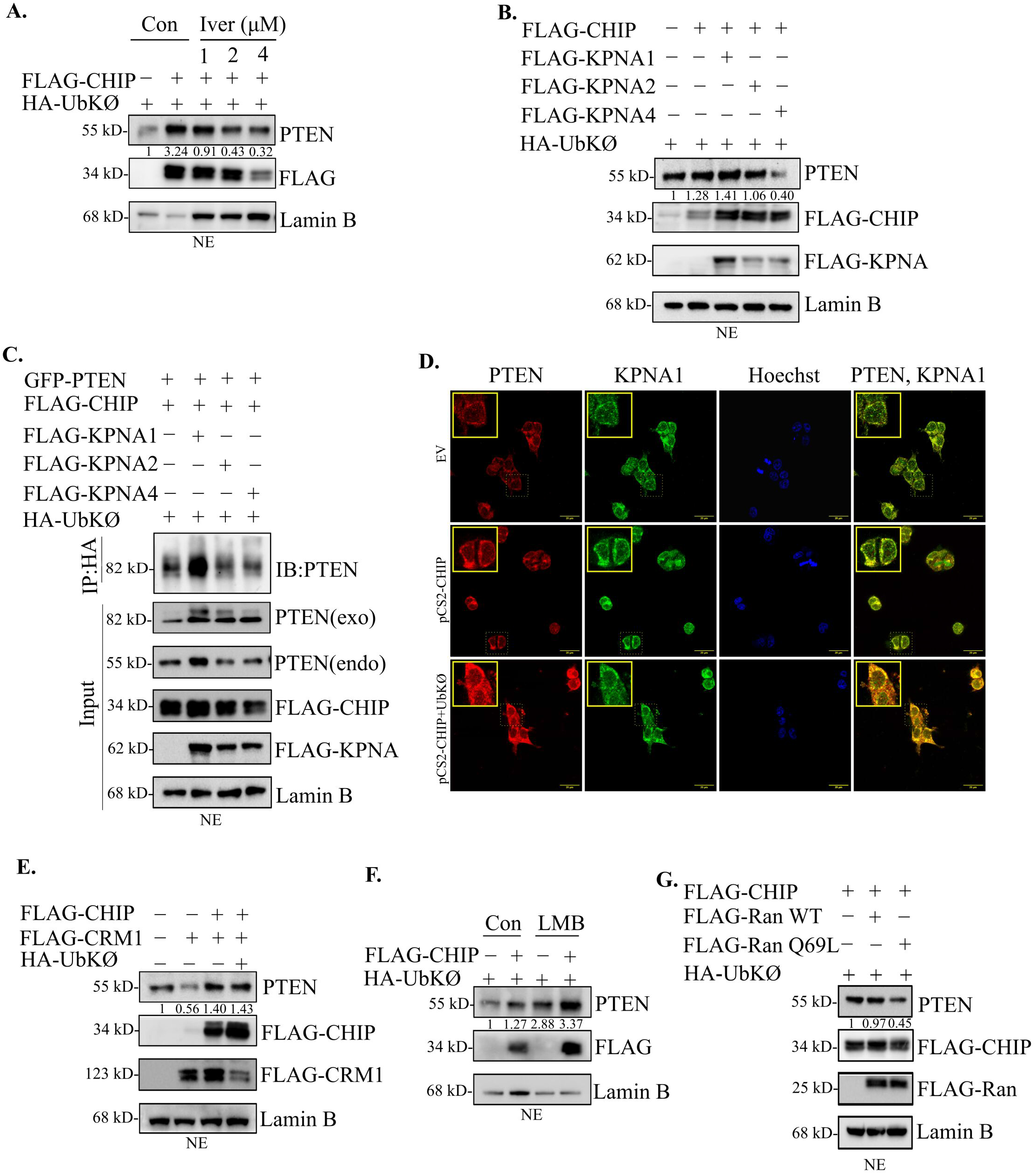
Nucleo-cytoplasmic shuttling of monoubiquitinated PTEN is dependent on nuclear import/export machinery. All experiments were performed in HEK293 cells. (**A**) Cells were transfected with FLAG-CHIP and HA-UbKØ and treated with increasing doses of Ivermectin. Nuclear lysates were probed for PTEN and FLAG (exogenous CHIP). (**B**) FLAG-CHIP and HA-UbKØ were overexpressed with either FLAG-KPNA1, FLAG-KPNA2, or FLAG-KPNA4. Nuclear lysates were immunoblotted for PTEN, FLAG-CHIP, and FLAG-KPNA. (**C**) GFP-PTEN, FLAG-CHIP, and HA-UbKØ were co-transfected with either FLAG-KPNA1, FLAG-KPNA2, or FLAG-KPNA4. Nuclear lysates were immunoprecipitated with anti-HA antibody and immunoblotted for PTEN. (**D**) EV, or pCS2-HA-CHIP, or pCS2-HA-CHIP and HA-UbKØ were co-transfected. Cells were stained with primary antibodies against PTEN or KPNA1, followed by secondary antibodies conjugated to AF594 (PTEN, red) or AF488 (KPNA1, green), and observed under fluorescent microscope at 60X magnification. Nuclei were visualized through Hoechst staining. (**E**) Cells were transfected with FLAG-CRM1 alone, or with FLAG-CHIP, or FLAG-CHIP and HA-UbKØ. Nuclear extracts were probed for PTEN, FLAG-CHIP, and FLAG-CRM1. (**F**) Cells were transfected with HA-UbKØ in presence or absence of FLAG-CHIP. Following transfection, the cells were treated with Leptomycin B (2 ng/ml for 24 hrs). Nuclear extracts were probed for PTEN and FLAG (exogenous CHIP). (**G**) FLAG-CHIP and HA-UbKØ were co-transfected with FLAG-RanWT or FLAG-RanQ69L. Nuclear extracts were immunoblotted for PTEN, FLAG-CHIP, and FLAG-Ran. 1 µg of each DNA construct was used in transfection for overexpression of cloned gene.

CRM1 mediates Ran-dependent active nuclear export of PTEN.^6^ Expressing CRM1 alone led to the expected decrease in nuclear PTEN. However, co-expressing CHIP with CRM1 counteracted this effect, further aided by UbKØ co-expression (Fig. 3E). While CHIP-mediated monoubiquitination increased nuclear PTEN level in untreated HEK293 cells, treatment with Leptomycin B, a CRM1 inhibitor, further enhanced it (Fig. 3F). Co-expressing CHIP and UbKØ with or without wild type Ran protein resulted in similar nuclear PTEN level, which was disrupted when GTPase activity deficient Ran-Q69L mutant was used, suggesting the involvement of Ran (Fig. 3G). In summary, the results showed that the components of nuclear import/export machinery are required for the shuttling of monoubiquitinated PTEN.

### CHIP and PTEN interact within the nuclear compartment where nuclear CHIP exerts its E3 ligase activity on nuclear PTEN

The presence of CHIP inside the nuclear compartment is well-established and various substrates of nuclear CHIP have been discovered.^19^ To detect whether PTEN was also among them, we performed a coimmunoprecipitation analysis from the nuclear fraction after co-expressing PTEN and CHIP with or without UbKØ. This revealed an interaction between exogenous CHIP and PTEN in both cases, although nuclear PTEN level was elevated under UbKØ-overexpressed condition (Fig. 4A). The reverse immunoprecipitation also indicated that nuclear PTEN interacted with both endogenous and exogenous CHIP, and the interaction increased upon UbKØ overexpression (Fig. 4B). Since PTEN lacks classical NLS, to ensure its nuclear localization, we used an NLS-tagged PTEN, which also coimmunoprecipitated with endogenous CHIP (Fig. 4C). Furthermore, immunocytochemistry analysis corroborated that compared to the endogenous condition, NLS-PTEN exhibited increased nuclear localization and enhanced interaction with nuclear CHIP (Fig. 4D).

**Figure 4.**
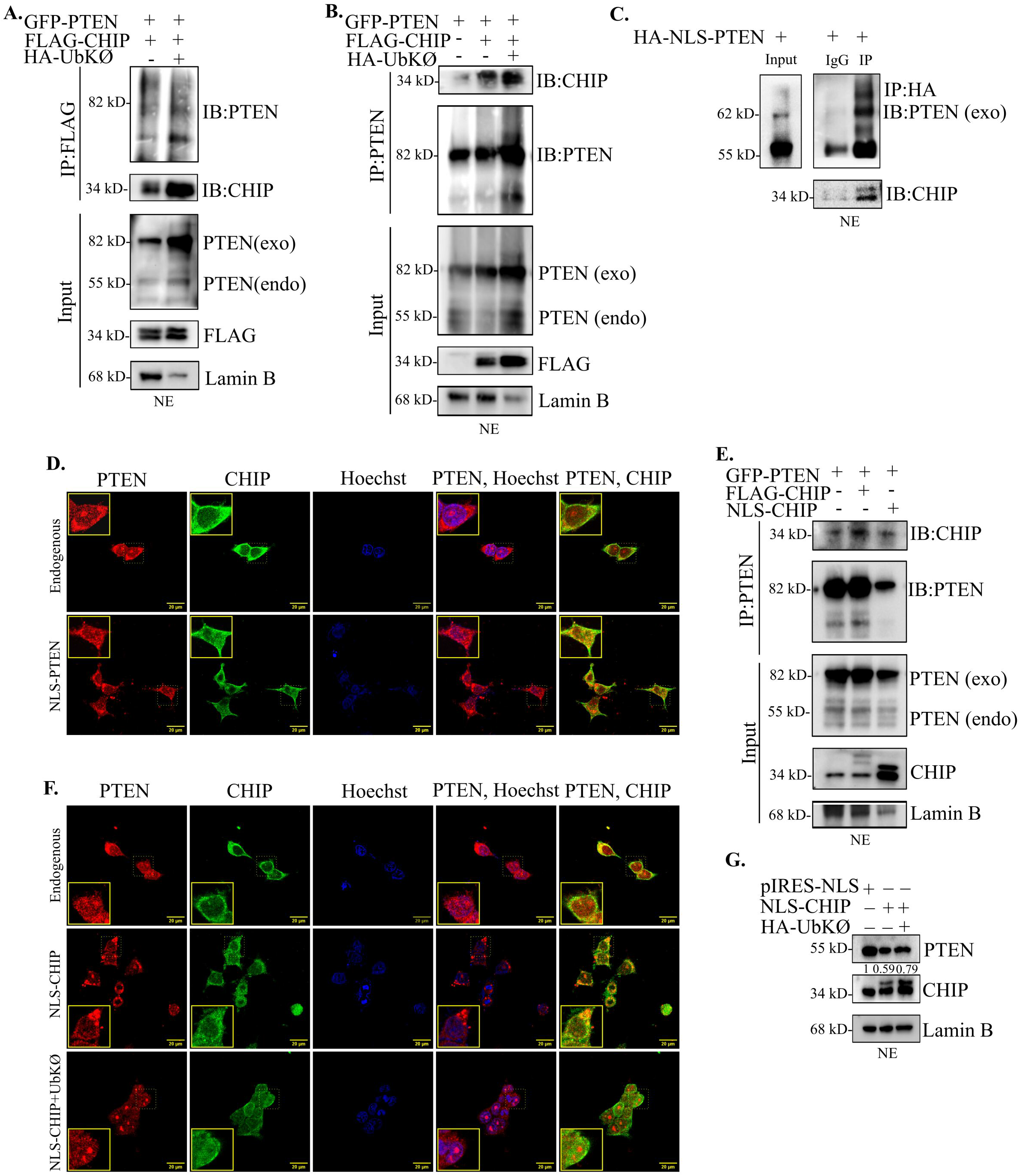
CHIP and PTEN interact within the nuclear compartment where nuclear CHIP exerts its E3 ligase activity on nuclear PTEN. All experiments were performed in HEK293 cells. (**A**) GFP-PTEN and FLAG-CHIP were co-transfected with or without HA-UbKØ. Nuclear extracts were immunoprecipitated with anti-FLAG antibody and immunoblotted for PTEN and CHIP. (**B**) Cells were transfected with GFP-PTEN alone, or GFP-PTEN and FLAG-CHIP, with or without HA-UbKØ. Nuclear extracts were immunoprecipitated with anti-PTEN and probed for CHIP and PTEN. (**C**) HA-NLS-PTEN was overexpressed. After immunoprecipitating the nuclear extract with anti-HA antibody, they were immunoblotted for PTEN and CHIP. (**D**) Either untransfected or HA-NLS-PTEN transfected cells were stained with primary antibodies against PTEN and CHIP, followed by secondary antibodies conjugated to AF594 (PTEN, red) or AF488 (CHIP, green). (**E**) GFP-PTEN was transfected either alone, or with FLAG-CHIP or NLS-CHIP. Nuclear lysates were immunoprecipitated with anti-PTEN antibodies and immunoblotted for CHIP and PTEN. (**F**) Untransfected, only NLS-CHIP transfected, or NLS-CHIP and HA-UbKØ transfected cells were stained with primary antibodies against PTEN and CHIP, followed by secondary antibodies conjugated to AF594 (PTEN, red) or AF488 (CHIP, green). (**G**) Cells were transfected with either pIRES-NLS, or NLS-CHIP, or NLS-CHIP and HA-UbKØ. Nuclear extracts were probed for PTEN and CHIP. 1 µg of each DNA construct was used in transfection for overexpression of cloned gene. For immunofluorescence microscopy, 60X magnification was used. Nuclei were visualized through Hoechst staining.

Next, we wanted to see the effects when, instead of PTEN, CHIP was forced into the nucleus. Upon transfecting GFP-PTEN in HEK293 cells alone, with CHIP, or with an NLS-tagged CHIP, we found that PTEN interacted with both endogenous and exogenous CHIP as well as NLS-CHIP. However, the forced nuclear localization of CHIP diminished nuclear PTEN level (Fig. 4E), which was further confirmed by immunocytochemical analysis (Fig. 4F). This implied a possible role of nuclear CHIP in driving degradation of nuclear PTEN. Western blot analysis from the nuclear lysates revealed similar results (Fig. 4G). However, co-expression of UbKØ with NLS-CHIP could reverse this effect and partially restore nuclear PTEN (Fig. 4F-G).

These experiments collectively demonstrate an interaction between CHIP and PTEN inside the nuclear compartment, facilitated by monoubiquitination. Nuclear CHIP acts as a polyubiquitinating E3 ligase of nuclear PTEN while still being able to monoubiquitinate it.

### CHIP executes nuclear translocation of PTEN in conjunction with other E3 ligases and deubiquitinases and requires both enzymatic and co-chaperone activities

Next, we wanted to see how CHIP acted alongside other E3 ligases and deubiquitinases implicated in PTEN nucleo-cytoplasmic shuttling. Both CHIP and XIAP were found to play a significant role in maintaining the nuclear pool of endogenous (Fig. 5A) and exogenous (Fig. S2A) PTEN. Consequently, CHIP exhibited a predominant role in maintaining the stability of cellular PTEN, followed by XIAP (Fig. S2B). NEDD4-1 increased exogenous nuclear and total PTEN levels, but not endogenous nuclear fraction. In all cases, UbKØ was co-expressed to enhance monoubiquitination by these bona fide E3 ligases.

**Figure 5.**
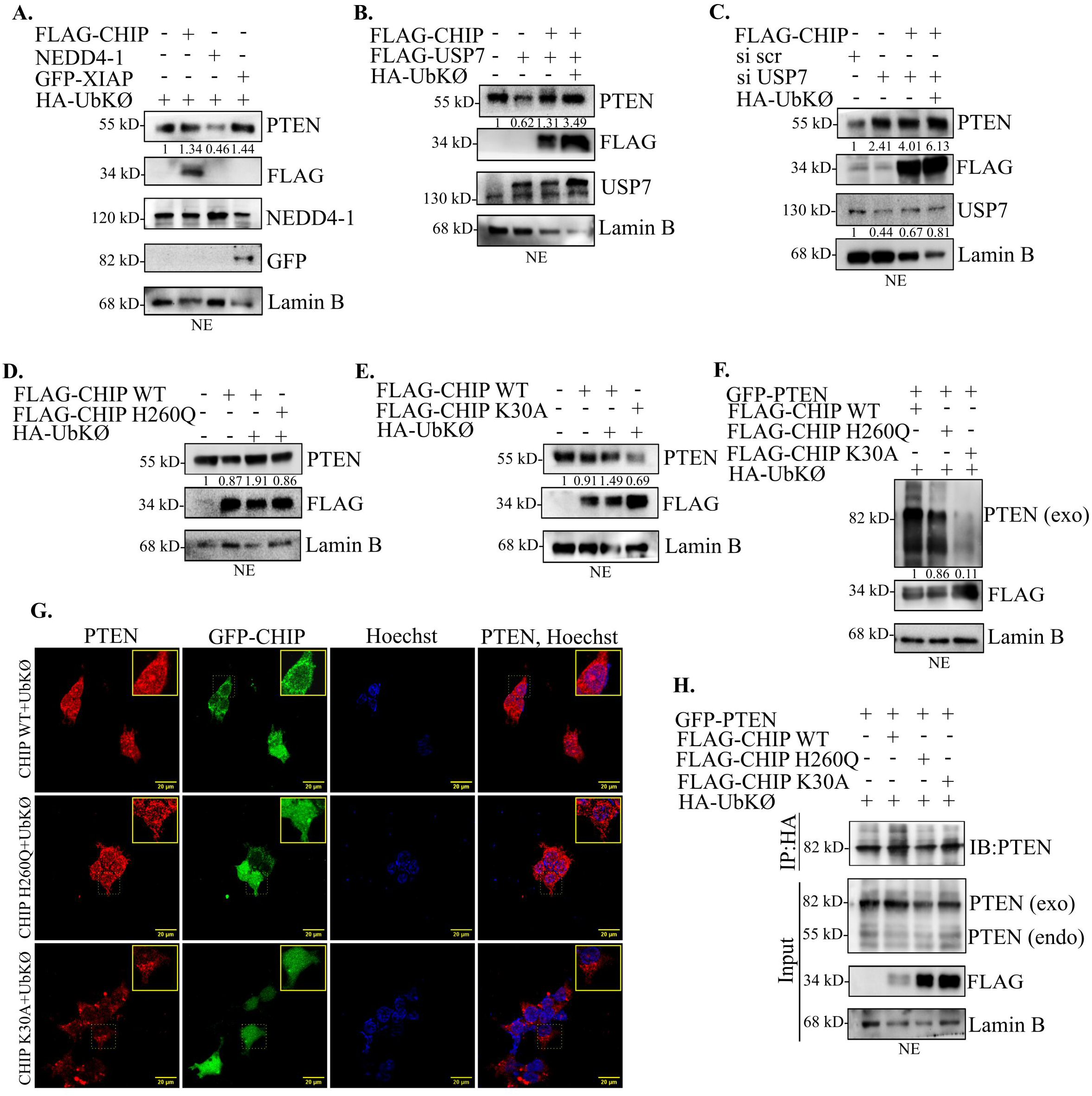
CHIP executes nuclear translocation of PTEN in conjunction with other E3 ligases and deubiquitinases and requires both enzymatic and co-chaperone activities. All experiments were performed in HEK293 cells. (**A**) Cells were transfected with HA-UbKØ and either FLAG-CHIP, or HA-NEDD4-1, or GFP-XIAP. Nuclear extracts were probed for PTEN, FLAG (exogenous CHIP), NEDD4-1, and GFP (exogenous XIAP). (**B**) FLAG-USP7 was transfected either alone or with FLAG-CHIP or with FLAG-CHIP and HA-UbKØ. Nuclear lysates were immunoblotted for PTEN, FLAG (exogenous CHIP), and USP7. (**C**) Cells were transfected with either scrambled siRNA or USP7 siRNA, alone or with FLAG-CHIP or FLAG-CHIP and HA-UbKØ. Nuclear extracts were probed for PTEN, USP7, and FLAG (exogenous CHIP). (**D**) Cells were transfected with FLAG-CHIP alone, or FLAG-CHIP and HA-UbKØ, or FLAG-CHIP-H260Q and HA-UbKØ. Nuclear extracts were probed for PTEN and FLAG (exogenous CHIP). (**E**) Cells were transfected with FLAG-CHIP alone, or FLAG-CHIP and HA-UbKØ, or FLAG-CHIP-K30A and HA-UbKØ. Nuclear extracts were probed for PTEN and FLAG (exogenous CHIP). (**F**) GFP-PTEN and Ha-UbKØ were co-transfected with either FLAG-CHIP, or FLAG-CHIP-H260Q, or FLAG-CHIP-K30A. Nuclear extracts were probed for PTEN and FLAG (exogenous CHIP). (**G**) Cells were transfected with HA-UbKØ and FLAG-CHIP, or FLAG-CHIP-H260Q, or FLAG-CHIP-K30A, stained with primary antibodies against PTEN, followed by secondary antibodies conjugated to AF594 (PTEN, red) and observed under fluorescent microscope at 60X magnification. GFP was observed to determine CHIP (wild type or mutants) overexpression. Nuclei were visualized through Hoechst staining. (**H**) GFP-PTEN and Ha-UbKØ were co-transfected with FLAG-CHIP, or FLAG-CHIP-H260Q, or FLAG-CHIP-K30A. Nuclear extracts were immunoprecipitated with anti-HA antibody and immunoblotted with PTEN. 1 µg of each DNA construct was used in transfection for overexpression of cloned gene.

The overexpression of USP7, a deubiquitinase involved in PTEN nuclear egress,^37^ resulted in a decrease in PTEN nuclear fraction, although co-expression of CHIP was able to restore nuclear PTEN level, UbKØ overexpression further enhancing this effect (Fig. 5B). In contrast, siRNA-mediated depletion of USP7 led to an increase in the nuclear PTEN, which was further stabilized by co-transfection of CHIP, and even more stabilized when UbKØ was co-expressed (Fig. 5C).

Further, CHIP-H260Q, the enzymatic mutant of CHIP, failed to increase the nuclear fraction of PTEN even during forced monoubiquitination, suggesting CHIP’s enzymatic activity is required for PTEN nuclear import (Fig. 5D). Consequently, CHIP-H260Q could not restore the total cellular level of PTEN, either (Fig. S2C). Using CHIP-K30A, a co-chaperone activity-deficient mutant of CHIP, we found that CHIP-WT but not CHIP-K30A could increase the level of nuclear (Fig. 5E) and total cellular (Fig. S2D) PTEN, even under monoubiquitination-promoting conditions, indicating that CHIP-mediated nuclear import and stabilization of PTEN occur in a chaperone-dependent way. Both enzymatic and co-chaperone activities were indispensable for the nuclear import of exogenous PTEN as well (Fig. 5F). This was further supported by immunocytochemical analysis, where, as compared to CHIP-WT, PTEN nuclear distribution decreased with CHIP-H260Q and CHIP-K30A (Fig. 5G). An immunoprecipitation experiment also confirmed that monoubiquitinated nuclear PTEN level was reduced when the mutants were used (Fig. 5H).

Taken together, CHIP acts alongside other E3 ligases and deubiquitinases to regulate nuclear distribution of PTEN, using both its enzymatic and co-chaperone activities.

### CHIP-mediated nuclear import of monoubiquitinated PTEN depends on K13 and K289 residues of PTEN

Next, we wanted to determine whether K13 and K289 residues, the predominant sites for PTEN’s monoubiquitination-mediated nuclear translocation^13,38,39^ were involved here. For this, we used PTEN-UbDM (Ubiquitin double mutant, PTEN-K13R/K289E), a PTEN mutant with both K13 and K289 residues mutated to arginine and glutamic acid, respectively, thus rendering them incapable of ubiquitination. When co-transfected with CHIP and UbKØ in HEK293 cells, wild type PTEN (PTEN-WT) but not PTEN-UbDM showed increased nuclear localisation (Fig. 6A). Coimmunoprecipitation analysis from the nuclear fraction indicated that the PTEN-UbDM interaction with exogenous CHIP was lower than that of PTEN-WT. The cytoplasmic fraction, however, yielded the opposite result (Fig. 6B). When PTEN-WT and PTEN-UbDM were co-transfected with CHIP and either UbWT or UbKØ, PTEN-WT but not PTEN-UbDM revealed distinct monoUb adduct and enhanced protein expression under UbKØ co-expression (Fig. 6C).

**Figure 6.**
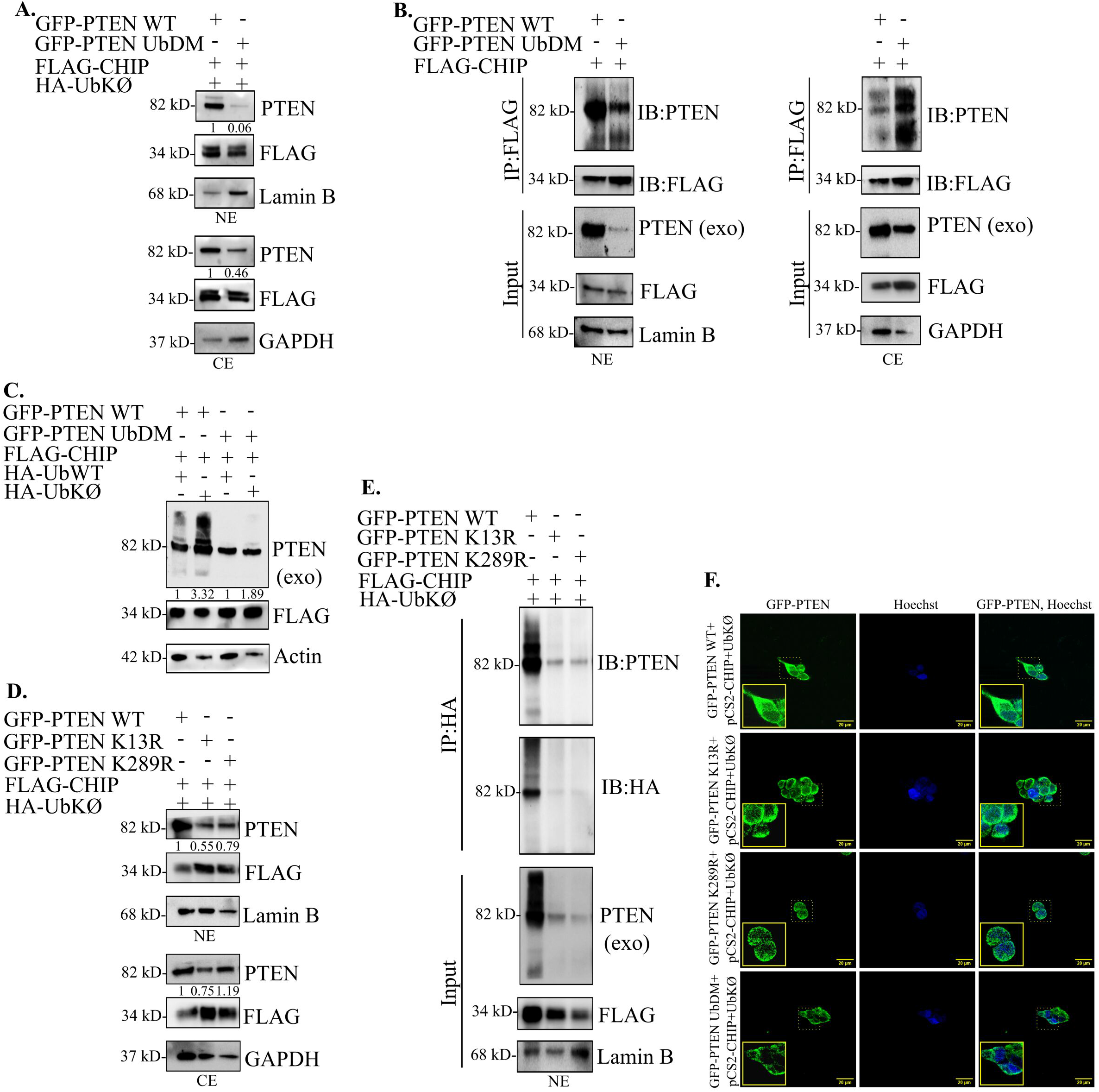
CHIP-mediated nuclear import of monoubiquitinated PTEN depends on K13 and K289 residues of PTEN. All experiments were performed in HEK293 cells. (**A**) FLAG-CHIP and HA-UbKØ were co-transfected with either GFP-PTEN-WT or GFP-PTEN-UbDM. Nuclear and cytoplasmic extracts were probed for PTEN and FLAG (exogenous CHIP). (**B**) Cells were transfected with FLAG-CHIP and either GFP-PTEN-WT or GFP-PTEN-UbDM. Nuclear and cytoplasmic extracts were immunoprecipitated with anti-FLAG antibody and immunoblotted for PTEN and FLAG (exogenous CHIP). (**C**) GFP-PTEN-WT or GFP-PTEN-UbDM was co-transfected with FLAG-CHIP and either HA-UbWT or HA-UbKØ. Whole cell lysates were probed for PTEN and FLAG (exogenous CHIP). (**D**) FLAG-CHIP and HA-UbKØ were co-transfected with either GFP-PTEN-WT, GFP-PTEN-K13R, or GFP-PTEN-K289R. Nuclear and cytoplasmic extracts were probed for PTEN and FLAG (exogenous CHIP). (**E**) FLAG-CHIP and HA-UbKØ were co-transfected with either GFP-PTEN-WT, GFP-PTEN-K13R, or GFP-PTEN-K289R. Nuclear extracts were immunoprecipitated with anti-HA antibody and immunoblotted for PTEN and HA. (**F**) pCS2-HA-CHIP and HA-UbKØ were co-transfected with GFP-PTEN-WT, GFP-PTEN-K13R, GFP-PTEN-K289R, or GFP-PTEN-UbDM. The localization of GFP-tagged PTEN (wild type or mutant) was observed under fluorescent microscope at 60X magnification. Nuclei were visualized through Hoechst staining. 1 µg of each DNA construct was used in transfection for overexpression of cloned gene.

Moreover, co-transfection of monoubiquitin mutants PTEN-K13R and PTEN-K289R with CHIP and HA-UbKØ showed diminished nuclear localization of both, as compared to PTEN-WT (Fig. 6D). Co-immunoprecipitation experiment from the nuclear fraction revealed monoubiquitinated forms of PTEN-WT but not PTEN-K13R or PTEN-K289R (Fig. 6E). Immunocytochemical analysis further confirmed that nuclear level of PTEN-WT was greater than those of PTEN-K13R, PTEN-K289R, and PTEN-UbDM (Fig. 6F). In summary, the K13 and K289 residues of PTEN were required for its CHIP-mediated monoubiquitination and subsequent nuclear import.

### CHIP-mediated nuclear import of PTEN augments its tumor suppressive functions

Next, we wanted to assess whether CHIP-mediated import could enhance the established tumor suppressive abilities of nuclear PTEN, such as increasing apoptosis,^6,7^ decreasing cell viability and cell proliferation.^40^

Upon co-expressing CHIP and UbKØ in HEK293, HCT 116, HepG2, and HeLa cells, we detected increased cellular levels of cleaved PARP, cleaved Caspase 3, and pro-apoptotic Bax and Bim as compared to the control, indicating higher apoptosis (Fig. 7A). To determine whether CHIP-mediated monoubiquitination of PTEN was responsible for this, an shRNA against PTEN (Fig. S3A) was co-transfected. PTEN knockdown decreased the apoptotic markers, counteracting the effects of CHIP and UbKØ. MTT assays in HEK293 and HCT 116 cells indicated that monoubiquitination by CHIP significantly diminished the cell viability, while the addition of sh PTEN nullified the effects by further increasing viability (Fig. 7B). Wound healing experiments from HEK293, HCT 116, HepG2, and HeLa cells revealed that, compared to the control, CHIP co-expression with UbKØ resulted in diminished wound healing. However, sh PTEN co-expression reversed this effect, leading to significantly increased cell proliferation and wound healing (Fig. 7C). In colony formation assays using HEK293, HCT 116, HepG2, and HeLa cells, CHIP-mediated monoubiquitination led to fewer colonies, an effect significantly abrogated by sh PTEN co-expression (Fig. 7D). Collectively, all these experiments indicate that CHIP can result in decreased cell viability, cell proliferation, and colony formation ability, as well as enhanced apoptotic potential, by monoubiquitinating and stabilizing PTEN. That PTEN knockdown could effectively abolish these outcomes further proved that CHIP mediates these tumor suppressive functions through PTEN.

**Figure 7.**
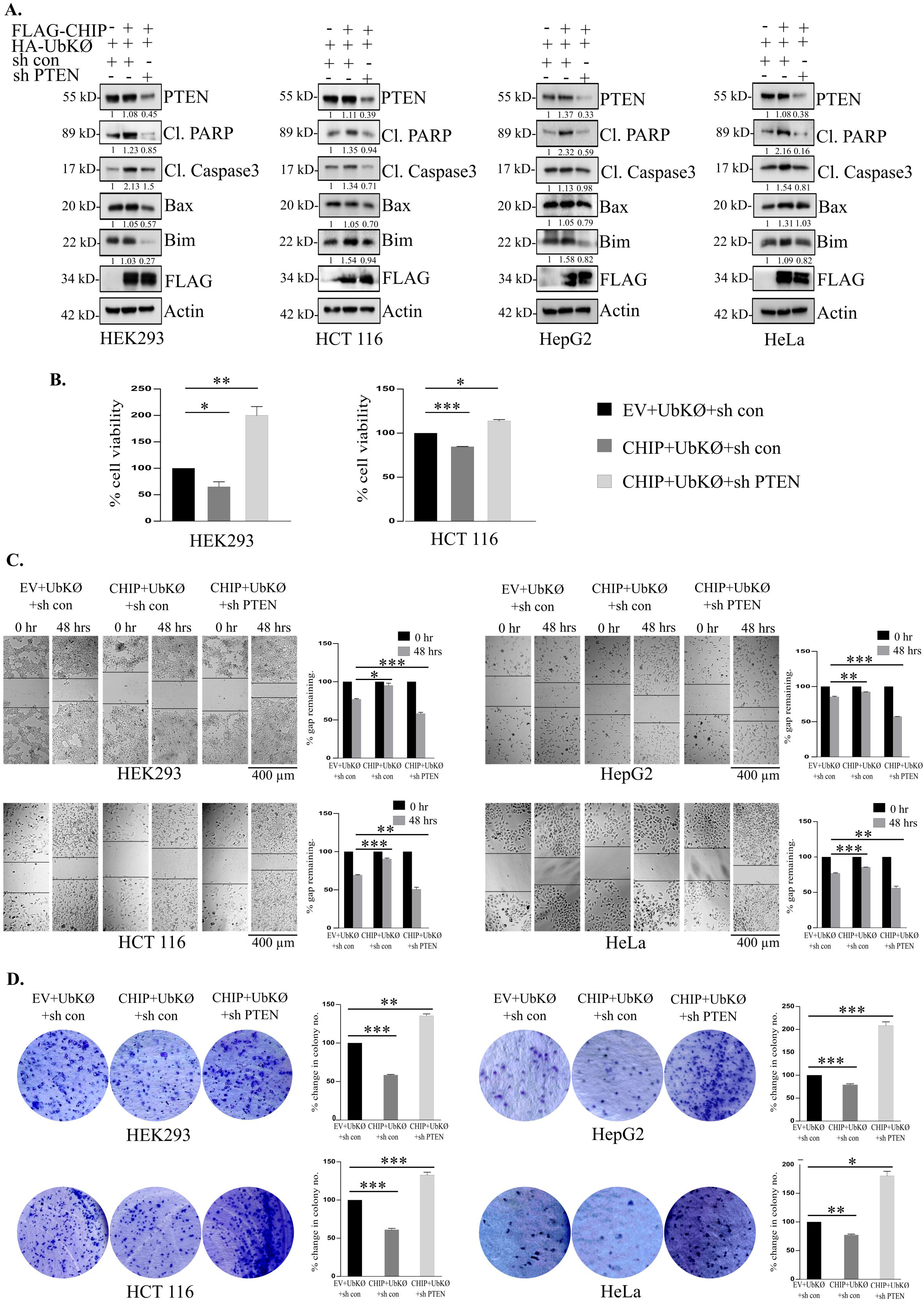
CHIP-mediated nuclear import of PTEN augments its tumor suppressive functions. (**A**) HEK293, HCT 116, HepG2, and HeLa cells were transfected with FLAG-CHIP alone or FLAG-CHIP and HA-UbKØ, with either sh control or sh PTEN. Whole cell lysates were probed for PTEN, Cleaved PARP, Cleaved caspase 3, Bax, Bim, and FLAG. (**B**) HEK293 and HCT 116 cells were subjected to MTT assay following overexpression of FLAG-CHIP alone or FLAG-CHIP and HA-UbKØ, with either sh control or sh PTEN. The average absorbance values of three biological repeats were plotted as percent change in cell viability. (**C**) HEK293, HCT 116, HepG2, and HeLa cells were transfected with FLAG-CHIP alone or FLAG-CHIP and HA-UbKØ, with either sh control or sh PTEN. After making transverse scratches on the plates with 10ul sterile pipette tips, the images were captured at the specified time points. Scale bar-400µm. Bar diagrams represent the percentage of gap remaining averaged from three biological repeats. (**D**) HEK293, HCT 116, HepG2, and HeLa cells were transfected with FLAG-CHIP alone or FLAG-CHIP and HA-UbKØ, with either sh control or sh PTEN. After 15 days, the cell colonies were stained with crystal violet. Bar diagrams represent the percent change in colony numbers averaged from three biological repeats. 1 µg of each DNA construct was used in transfection for overexpression of cloned gene. Error bars in all the indicated subfigures represent mean (+) s.d. from three independent biological repeats. Indicated *P* values were calculated using Student’s *t*-test, and represented as *\* (P* ≤ 0.05), ** (*P* ≤ 0.01) and *** (*P* ≤ 0.001).

### CHIP-induced nuclear translocation modulates transcriptional activities of PTEN

The enhanced tumor suppressive functions of PTEN following CHIP-mediated monoubiquitination made us to explore possible alterations in the transcriptional activities of nuclear PTEN. We selected Rad51 and Dre1 as target genes upregulated by PTEN and VEGF as a target gene downregulated by PTEN.^20^ cBioPortal analysis^25^ in colon, liver, and cervical cancers from TCGA PanCancer dataset and Cancer Cell Line Encyclopedia datasets revealed a positive correlation between PTEN and Rad51 genes in colon cancer but not in liver cancer, while PTEN and Dre1 genes showed a positive correlation in both. PTEN and VEGF exhibited a negative correlation in all three cancers. Significant correlations of PTEN with Rad51 and Dre1 were lacking in cervical cancer (Fig. 8A). However, Cancer Cell Line Encyclopedia data showed a positive correlation between PTEN-Rad51 and PTEN-Dre1, while PTEN-VEGF showed a negative correlation (Fig. 8B). Next, we transfected HEK293 cells with UbKØ, with or without CHIP. qPCR analysis revealed significantly elevated levels of Rad51 and Dre1 mRNAs, as well as reduced VEGF mRNAs upon co-expressing CHIP with UbKØ (Fig. 8C), suggesting CHIP-mediated monoubiquitination enhanced transcriptional upregulation of Rad51 and Dre1 and downregulation of VEGF by PTEN.

**Figure 8.**
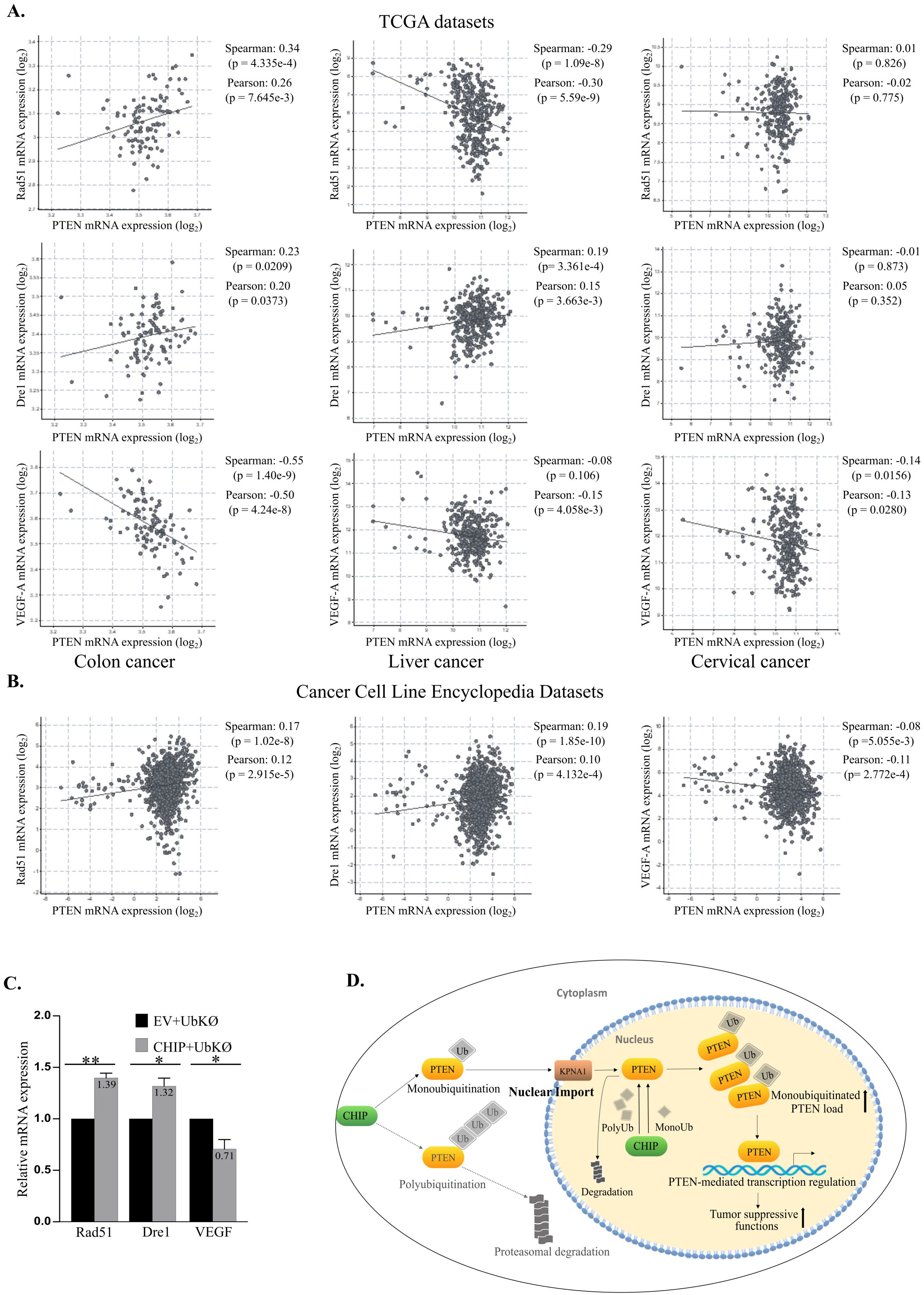
CHIP-induced nuclear translocation modulates transcriptional activities of PTEN. (**A**) cBioPortal analysis of the correlation between PTEN-Rad51 (upper row), PTEN-Dre1 (middle row), and PTEN-VEGF (lower row) in colon, liver, and cervical cancers (*P* < 0.05) from TCGA PanCancer datasets. (**B**) cBioPortal analysis of correlation between PTEN-Rad51, PTEN-Dre1, and PTEN-VEGF (*P* < 0.05) from Cancer Cell Line Encyclopedia datasets. (**C**) HA-UbKØ was co-transfected in HEK293 cells with either EV or FLAG-CHIP. cDNA was synthesized and qPCR performed to check the mRNA levels of Rad51, Dre1, and VEGF. Error bars in all the indicated subfigures represent mean (+) s.d. from three independent biological repeats. Indicated *P* values were calculated using Student’s *t*-test, and represented as *\* (P* ≤ 0.05), ** (*P* ≤ 0.01) and *** (*P* ≤ 0.001). (**D**) Proposed model: A schematic representation of CHIP-induced monoubiquitination of PTEN followed by nuclear transport and enhanced tumor suppression.

## Discussion

Nuclear PTEN has been implicated in a plethora of tumor-suppressive activities, complementing the functions of its cytoplasmic counterpart. However, different cancers show considerably variable nuclear PTEN levels. While the correlation between nuclear PTEN loss and disease progression is distinct in colorectal cancer,^28–32^ the reports from cervical^33,34^ and hepatic^35,36^ cancers are somewhat elusive. The mechanisms behind this differential localization and maintenance of nuclear PTEN also remain unclear. In this study, we sought to explore the probable non-canonical monoubiquitination of PTEN by chaperone-associated E3 ligase CHIP, followed by the former’s nuclear import.

We evaluated PTEN mRNA and protein expressions using multiple bioinformatic tools in colon, liver, and cervical cancer datasets as well as the selected cell lines as background study. While delving deeper into the mechanisms, the increased cellular PTEN level under monoubiquitinating conditions in all cell lines suggested that CHIP, through its non-canonical activity, could protect PTEN by diverting it from the degradation pathway. Further, immunoprecipitation of ubiquitinated PTEN revealed not merely a polyubiquitin smear but distinct, lower molecular weight monoubiquitinated bands, confirming non-canonical multi-monoubiquitination by CHIP. Additionally, A closer look at cytonuclear distribution of PTEN further indicated that CHIP-mediated monoubiquitination could facilitate PTEN’s nuclear import. The observed rise in nuclear PTEN under monoubiquitination-promoting conditions was not an indirect effect of hindering PTEN degradation but a direct result of monoubiquitination. Reduced nuclear level of PTEN upon CHIP knockdown has been previously described, which could not be restored even after introducing UbKØ in the system, highlighting the importance of CHIP as a monoubiquitinating E3 ligase. Incidentally, these findings partially resemble a previous study, where XIAP knockdown decreased PTEN poly- and monoubiquitination, the latter causing a decline in the nuclear fraction of PTEN, while overexpressed XIAP reduced total cellular PTEN by selectively promoting polyubiquitination. However, unlike XIAP overexpression, CHIP overexpression altered PTEN subcellular distribution by promoting its nuclear entry. Starting at almost 30 hrs post-transfection, this effect continued for at least 48 hrs. However, PTEN’s nuclear localization was much more pronounced with exogenous supply of monoubiquitination-promoting UbKØ.

The inhibition of import proteins led to diminished nuclear PTEN even after CHIP-mediated monoubiquitination, whereas inhibiting CRM1 caused increased accumulation of nuclear PTEN following monoubiquitination. We further found that Karyopherin α1 (Importin α5) is involved in driving monoubiquitinated PTEN into the nucleus. Although a previous report suggested that PTEN nuclear translocation did not involve direct binding with Importin α5, they nonetheless suggested an importin-mediated nuclear transport.^6^ Moreover, the study did not elaborate on the effects of post-translational modifications (PTMs) on nuclear entry. PTM such as NEDDylation can enhance PTEN’s interaction with specific importin proteins, including Importin α.^11^ Also, Importin11 can specifically recognize and drive nuclear translocation of monoubiquitinated PTEN.^16^ In this study, we propose a similar scenario where CHIP-mediated monoubiquitination leads to an increased affinity between PTEN and Karyopherin α1, with the latter facilitating the former’s nuclear import. Incidentally, our findings upheld the involvement of Ran GTPase in PTEN’s nuclear localization.

Although predominantly cytoplasmic, CHIP is also present in the nucleus, especially during cellular stress. Various nuclear substrates of CHIP have been discovered previously. However, our study reports for the first time an interaction between CHIP and PTEN inside the nucleus, with exogenous PTEN interacting with both endogenous and exogenous CHIP. UbKØ co-expression further augmented this interaction, possibly by making more PTEN available in the nucleus *via* increased import. The observed rise in PTEN-CHIP nuclear colocalization upon specifically targeting PTEN to the nucleus by using NLS-tagged PTEN further supported this claim. Surprisingly, when we forced CHIP into the nucleus by fusing an NLS, the nuclear-localized CHIP still interacted with yet yielded a reduced level of nuclear PTEN. This made us hypothesize a possible role of nuclear CHIP in destroying nuclear PTEN. So far, FBXO22 has been discovered as the sole E3 ligase specifically degrading nuclear PTEN *via* K48-mediated polyubiquitin linkage. Our findings place CHIP as yet another nuclear E3 ligase of PTEN. While CHIP-WT overexpression could aid PTEN’s nuclear entry albeit reducing its total cellular level, the accelerated degradation of nuclear PTEN by nuclear CHIP could be explained by the fact that nuclear pool of PTEN was more prone to degradation than cytoplasmic PTEN.^15^ However, under a monoubiquitination-inducing condition, this destructive role of NLS-CHIP was reverted, indicating that CHIP-mediated monoubiquitination was also possible within the nucleus.

We further sought to determine CHIP’s effects in relation to the other bona fide monoubiquitinating E3 ligases and deubiquitinases of PTEN. Although a former study reported that overexpressed XIAP did not affect PTEN subcellular localization,^14^ we found that co-expression of UbKØ and XIAP could raise nuclear and total cellular PTEN levels. While CHIP’s functional activity seemed at par with XIAP, we failed to detect these protective effects from NEDD4-1, despite the latter being the first-described E3 ligase with dual poly- and monoubiquitinating role. No alterations in PTEN subcellular localizations in the absence of NEDD4-1 have been reported,^41^ while another group observed that NEDD4-1 knockdown led to increased cytoplasmic PTEN but had no effects on nuclear PTEN.^15^ Hence, the regulatory role of NEDD4-1 could be extremely cell and tissue-specific and was not apparent here. Our findings corroborated USP7’s previously described role in causing PTEN’s nuclear exclusion,^37^ while also suggesting that CHIP co-expression could more than compensate for these effects. Although CHIP could drive nuclear translocation of its substrates^18^ by utilizing its intrinsic chaperone activity,^42^ our results indicated that along with the enzymatic activity, co-chaperone activity of CHIP was also required for its monoubiquitinating function. This corroborated our previous findings that CHIP’s interaction with and regulation of PTEN involved both the catalytic Ubox and chaperone-associated TPR domains. CHIP-mediated monoubiquitination of insulin receptor likewise involved both.^17^

In line with previous findings, our results involving two single-site mutants and a dual-site mutant confirmed the importance of K13 and K289 residues in CHIP-mediated monoubiquitination and nuclear localization. Further, various functional assays confirmed that while CHIP-mediated monoubiquitination increased PTEN level and exhibited a greater tumor suppressive potential by promoting apoptosis, decreasing cell viability, cell proliferation, and colony formation, sh RNA-mediated knockdown of endogenous PTEN abolished these effects. The accelerated protooncogenic outcomes in PTEN’s absence underscored the importance of maintaining this tumor suppressor at an adequate level. The significant alteration detected in PTEN’s transcriptional activities, due to its increased nuclear import upon monoubiquitination by CHIP, could further contribute to these enhanced tumor suppressive functions. A schematic representation of CHIP-induced monoubiquitination and subsequent nuclear import of PTEN has been depicted in Fig. 8D.

To the best of our knowledge, this is the first ever report exploring the non-canonical monoubiquitination of PTEN by its bona fide E3 ligase CHIP while also demonstrating their interactions within the nuclear compartment. The similar results obtained using either HEK293 cell line or cell lines from three different cancers suggested that this could be a general phenomenon and not specific to a particular cancer. The starkly opposing outcomes of monoubiquitination *vs* polyubiquitination in regulating PTEN protein demand further explorations into the mechanistic details. Since the same E3 ligases could carry out two different modes of ubiquitination and the same two lysine residues were involved in both, which mechanism would prevail could depend on several contexts. From the fact that overexpression *vs* knockdown of E3 ligases could bring out different results, we could surmise that the prevalent mode of ubiquitination will depend on the ligases’ availability and subcellular distribution. The specific E2 conjugating enzymes could also determine the type of ubiquitination. Moreover, in the case of CHIP and PTEN, cellular stress might promote monoubiquitination since both proteins translocate to nucleus under stress, and PTEN monoubiquitination has been implicated in nuclear import under stress conditions.^38^ A proper understanding of the underlying factors would be pivotal in the maintenance of the tumor suppressor PTEN.

## Acknowledgement

Authors sincerely acknowledge Dr. Syed Feroj Ahmed and Ms. Satadeepa Kal (ex-students of Dr. Mrinal K Ghosh) for their technical help in developing this project and current findings.

## Abbreviations

PTEN: Phosphatase and Tensin Homolog Deleted on Chromosome TEN
KPNA1: Karyopherin α1
CHIP: Carboxy terminus of Hsc70 Interacting Protein
NEDD4-1: Neural precursor cell Expressed Developmentally Down-regulated protein
XIAP: X-linked Inhibitor of Apoptosis protein.

## Supplementary data

Supplementary figures S1 – S3; List of all primers used.

