## Supplemental Data 1 for "E3 Ubiquitin Ligase CHIP Drives Monoubiquitination-mediated Nuclear Import of Tumor Suppressor PTEN"

### Supplementary Information

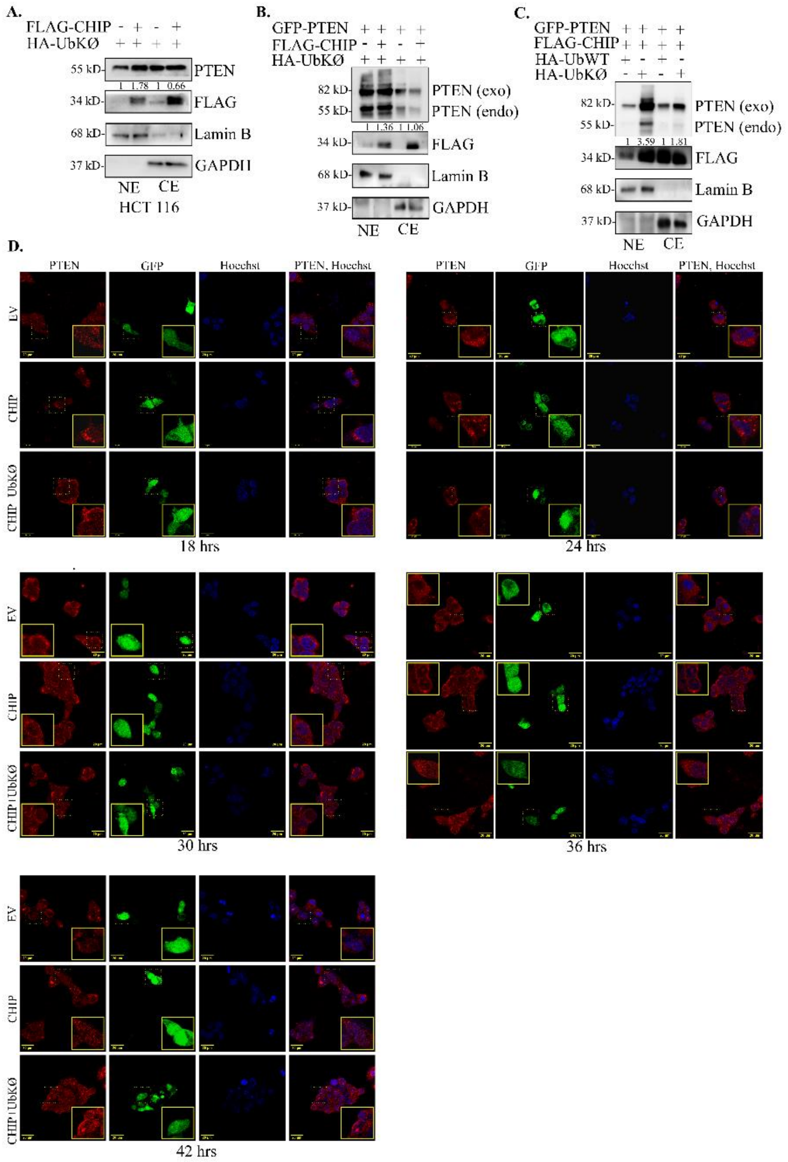


**Figure S1**

(A) HCT 116 cells were transfected with HA-UbKØ with or without FLAG-CHIP. Nuclear and cytoplasmic extracts were probed for PTEN and FLAG (exogenous CHIP). (B) HEK293 cells were transfected with GFP-PTEN and HA-UbKØ with or without FLAG-CHIP. Nuclear and cytoplasmic extracts were probed for PTEN and FLAG (exogenous CHIP). (C) HEK293 cells were transfected with GFP-PTEN and FLAG-CHIP with HA-UbWT or HA-UbKØ. Nuclear and cytoplasmic extracts were probed for PTEN and FLAG (exogenous CHIP). (D) HEK293 cells were transfected with either empty vector (EV), or FLAG-CHIP, or FLAG-CHIP and HA-UbKØ, EV and FLAG-CHIP GFP-tagged. 18 hrs, 24 hrs, 30 hrs, 36 hrs, and 42 hrs post-transfection, cells were fixed and stained with primary antibodies against PTEN, secondary antibodies conjugated to AF594 (PTEN, red), and observed under fluorescent microscope at 60X magnification. GFP was observed to determine EV or CHIP overexpression. Nuclei were visualized through Hoechst staining. 1 µg of each DNA construct was used in transfection for overexpression of cloned gene.

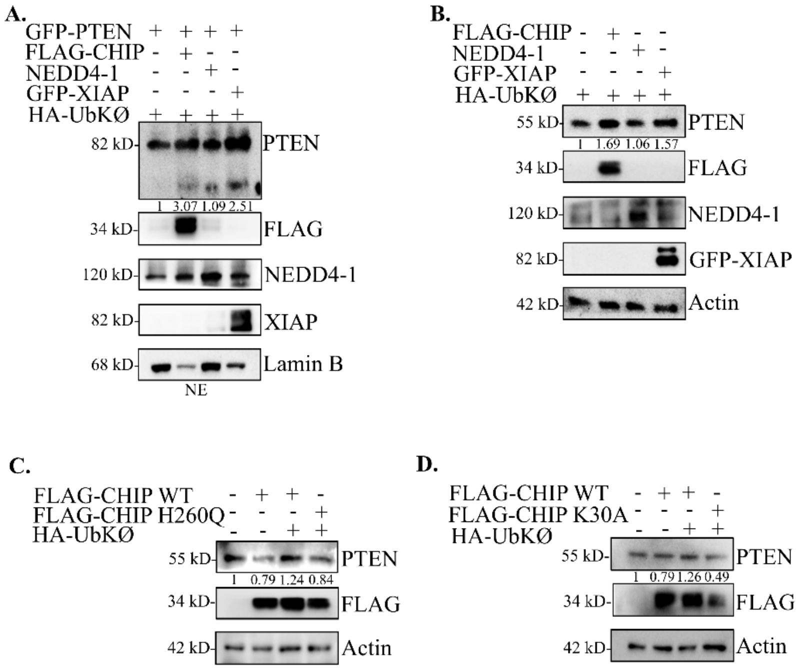


**Figure S2**

All experiments were performed in HEK293 cells. (A) Cells were transfected with GFP-PTEN and HA-UbKØ and either FLAG-CHIP, or HA-NEDD4-1, or GFP-XIAP. Nuclear extracts were probed for PTEN, FLAG (exogenous CHIP), NEDD4-1, and XIAP. (B) Cells were transfected with HA-UbKØ and either FLAG-CHIP, or HA-NEDD4-1, or GFP-XIAP. Whole cell lysates were probed for PTEN, FLAG (exogenous CHIP), NEDD4-1, and GFP (exogenous XIAP). (C) Cells were transfected with FLAG-CHIP alone, or FLAG-CHIP and HA-UbKØ, or FLAG-CHIP-H260Q and HA-UbKØ. Whole cell lysates were probed for PTEN and FLAG (exogenous CHIP). (D) Cells were transfected with FLAG-CHIP alone, or FLAG-CHIP and HA-UbKØ, or FLAG-CHIP-K30A and HA-UbKØ. Whole cell lysates were probed for PTEN and FLAG (exogenous CHIP). 1 μg of each DNA construct was used in transfection for overexpression of cloned gene.

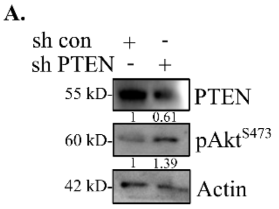


**Figure S3**

(A) HEK293 cells were transfected with either control shRNA or PTEN shRNA. The whole cell lysates were probed for PTEN and phospho-Akt. Actin was used as a loading control. 1 µg of each DNA construct was used in transfection for overexpression of cloned gene.

Supplementary Table 1

| | |
|---|---|
| PTENK13R | F: 5’-AGAGATCGTTAGCAGAAACAGAAGGAGATATCAAGAGGATG-3’ |
| | R: 5’-CATCCTCTTGATATCTCCTTCTGTTTCTGCTAACGATCTCT-3’ |
| PTENK289R | F: 5’-CAGGACCAGAGGAAACCTCAGAAAGGGTAGAAAATGGAAGTCTA-3’ |
| | R: 5’-TAGACTTCCATTTTCTACCCTTTCTGAGGTTTCCTCTGGTCCTG-3’ |

| | |
|---|---|
| 18s rRNA | F: 5’-GCTTAATTTGACTCAACACGGGC-3’ |
| | R: 5’-AGCTATCAATCTGTCAATCCTGTC-3’ |
| Rad51 | F: 5’-GGTCTGGTGGTCTGTGTTGA-3’ |
| | R: 5’-GGTGAAGGAAAGGCCATGTA-3’ |
| Dre1 | F: 5’-GAGTGCTACGATCCTGTAAC-3’ |
| | R: 5’-CATCACGGCTGTTGATTCTTC-3’ |
| VEGF | F: 5’-AGGAGGAGGGCAGAATCATCA-3’ |
| | R: 5’-CTCGATTGGATGGCAGTAGCT-3’ |

| | |
|---|---|
| sh PTEN | 5’-GGCACAAGAGGCCCTAGATT-3’ |
